# TwinEco: A Unified Framework for Dynamic Data-Driven Digital Twins in Ecology

**DOI:** 10.1101/2024.07.23.604592

**Authors:** Taimur Khan, Koen de Koning, Dag Endresen, Desalegn Chala, Erik Kusch

## Abstract

A Digital Twin (DT) is a virtual replica of a physical object or process that is continuously updated at a certain frequency and can steer change on the physical system, enabling a seamless integration of observation, understanding, and action. Although initially applied primarily in industry, DT is emerging as a powerful tool in ecology, offering new possibilities for dynamic simulations of change in the biosphere. However, since DTs are relatively new in this field, there is currently no standard framework to guide their conceptualisation and development. Thus, DTs in ecological applications are already experiencing fragmentation in software concepts and design philosophies, leading to incompatibilities across DT implementations. This fragmentation risks undermining the progress and potential of the DT concept in ecology. A unifying framework, such as TwinEco, can address these discrepancies and establish a cohesive foundation for the effective adoption and integration of DTs across ecological domains. TwinEco is a modular framework designed to aid and harmonise ecologists’ efforts to build DTs. In doing so, TwinEco focuses on three major design goals:

1. Modularity, flexibility, and interoperability of DTs facilitated through distinct “components” nested within DT “layers”.
2. Dynamic modeling of ecological processes and states that evolve over time.
3. Linking ecological modelling to downstream actions or decisions made on the ecological object or process of study.

TwinEco’s architecture builds upon the feedback loops and state management strategies introduced in the Dynamic Data-Driven Application Systems (DDDAS) paradigm, which has already inspired many DTs across scientific domains. We also discuss the usefulness and ease-of-use of TwinEco by demonstrating its applicability to computational case studies and suggesting future recommendations to the community of data infrastructure builders and modellers in regards to open considerations. By introducing a shared terminology and emphasising model-data fusion, TwinEco highlights the importance of a unified framework to avoid fragmentation in the burgeoning field of ecological digital twinning.

## 2. Introduction

In the rapidly evolving realm of environmental change and technology, just-in-time modelling and timely mitigational or adaptive actions become increasingly relevant. Ecological systems are dynamic, complex, and highly responsive to environmental factors, making them challenging to model and manage effectively (Carpenter and Brock, 2004; Donohue et al., 2016; Vermeiren et al., 2020). Understanding, predicting, effectively managing, and aligning human activities with these intricate systems are paramount for mitigating the profound impacts of human activities on the environment (Brown and Williams, 2015; Ruckelhaus et al., 2020; Newton, 2016). In response, ecological research has witnessed a remarkable evolution over the years, driven by technological advancements and the growing awareness of environmental issues. Nevertheless, much of contemporary ecological research informing management and policy continues to rely on models incorporating observational data at a given point in time without subsequent re-examination as additional data becomes available, thus producing static outputs (Zurell et al., 2021). While often informative, these traditionally static quantitative approaches fall short of capturing the dynamic, interconnected, and responsive nature of ecosystems (Damgaard, 2019). In response to these challenges, a transformative concept offering a promising shift to steer more informative quantification of ecosystem dynamics has emerged, the "digital twin”.

DT systems, rooted in the realm of cyber-physical systems, are virtual representations of physical entities or processes that capture real-time data and behaviour of system components, enabling dynamic simulation, monitoring, and analysis, while also providing feedback in the form of actions or decisions that directly influence the physical process or object under study. (Segovia & Garcia-Alfaro, 2022). Initially, these systems were designed for applications such as manufacturing, where they facilitate predictive maintenance, quality control, and process optimization (Wu & Li, 2021; Onaji et al., 2022). However, the potential for DT extends well beyond industry boundaries and is rapidly finding its place in the study and conservation of ecological systems (de Koning et al., 2023, Lecarpentier et al., 2024). In recognition of these opportunities, calls for proposals have been made across the European Union (European Commission, 2024a, European Commission, 2024b, CORDIS - European Commission, 2024), United Kingdom (UK Research and Innovation, 2024), and United States of America (NASA Technical Reports Server, 2024, NASA Earth Science Technology Office, 2024) to support development of a host of ecological DTs. Meanwhile, digital twinning is already a tool used for national and regional policy making in China (Tang & Chen, 2024, Brueck et al., 2024). Consequently, DTs are expected to rapidly establish themselves as a vital modelling approach to environmental and ecological research across geographic areas as well as sub-disciplines of ecology.

The necessary adaptation of twins from industry to ecological applications cannot be technically verbatim as ecological modelling presents its own unique challenges. Whereas traditional ecological modelling approaches often face limitations in their ability to update and adapt to changing conditions and respond to continuously evolving data on natural ecosystems, DTs are designed to iteratively update the process knowledge they create as well as the data products they incorporate and produce. Consequently, there is a growing recognition within the ecological research community that a paradigm shift is required to bridge the gap between observation, understanding, and action (Sharef et al., 2022). As the ecological modelling community explores avenues for overcoming said gap, modellers are increasingly exploring digital twinning as a possible solution (Lecarpentier et al., 2024).

Despite this pronounced interest in conceptualising a range of ecological DTs, digital twinning of ecological research objectives poses a number of problems requiring careful consideration. These correspond to considerations of <u>observation</u>, <u>understanding</u>, and <u>action/decision (Figure 1)</u> <u>(Lecarpentier et al., 2024)</u>. Firstly, incorporating observations into ecological modelling often suffers from considerable time lags. This is due to the workload involved in processing large volumes of heterogeneous data and the incomplete or delayed availability of critical datasets, compounded by inconsistent standards across ecological studies (Reichman et al., 2011). Secondly, the complexity of ecological systems poses a substantial obstacle. These systems involve nonlinear interactions, multi-scale processes, and stochastic variability, demanding advanced modelling approaches to capture their intricate dynamics accurately. Thirdly, translating insights from ecological DTs into actionable knowledge is complicated by the urgency of environmental changes and the practical and policy constraints associated with implementing direct interventions. Overcoming these challenges is essential to harness the full potential of ecological DTs for real-time monitoring, predictive modelling, and informed decision-making in environmental research and management.

**Figure 1:**
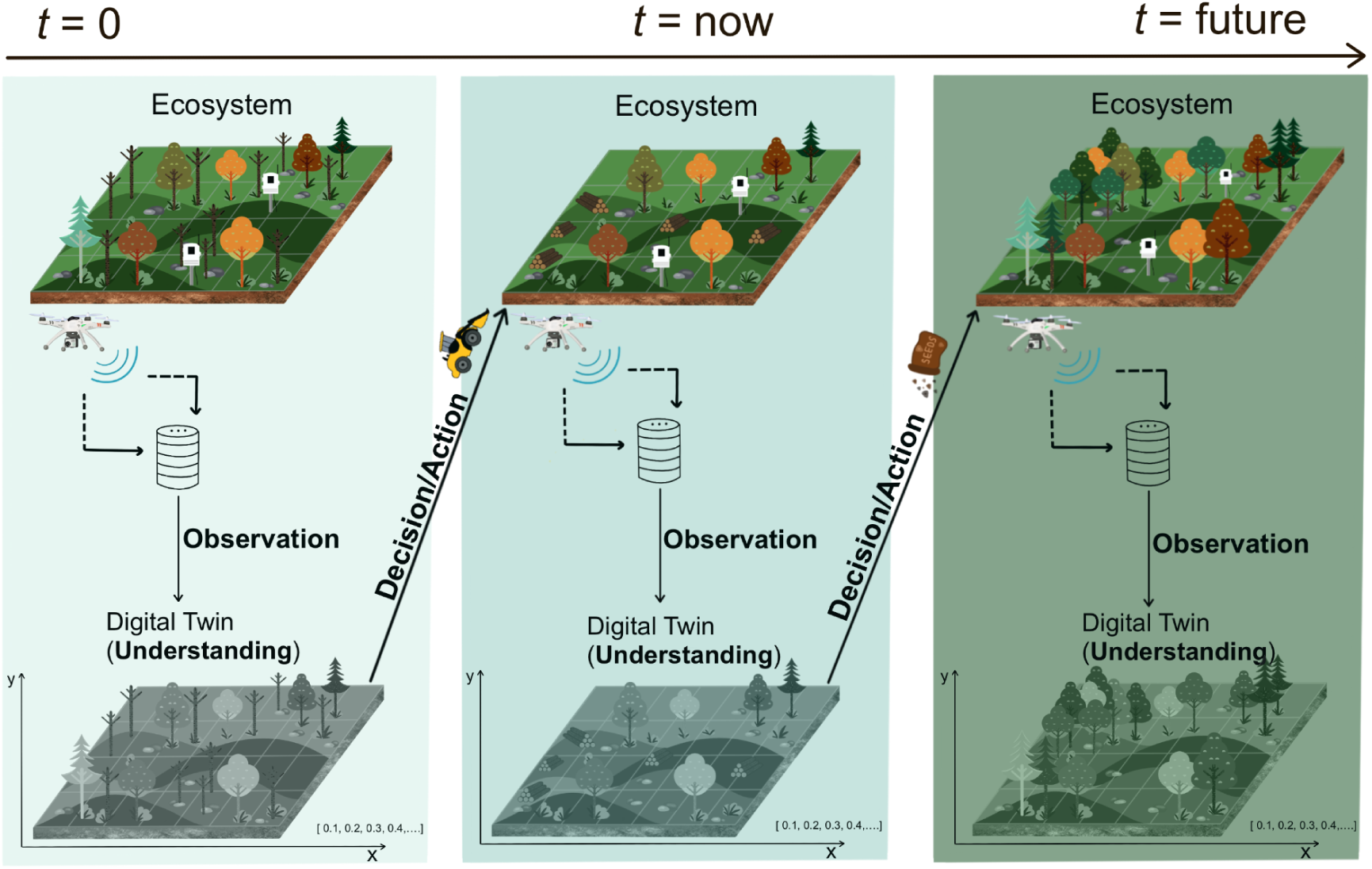
The dynamic integration of **observation, understanding, and decision/action** in a sample forestry Digital Twin (DT). At each timestep (t = 0, t = now, t = future), observational data from sensors and drones is ingested into the DT to update the state of the system and support real-time understanding of the forest. This process incorporates state space variables such as tree health, canopy cover, and soil moisture. The DT enables decision-making and control actions, such as planting trees or applying targeted soil nutrients, to achieve reward-driven ecological goals like enhanced biodiversity or carbon sequestration. Feedback loops between the ecosystem and the DT ensure adaptive management as conditions evolve over time. The iterative nature of this approach demonstrates the flexibility and modularity of the TwinEco framework in addressing complex ecological dynamics.

As the adoption of DT systems gains momentum in ecology, the associated complexities and challenges are becoming increasingly apparent across all disciplines of ecological research (Trantas et al., 2023; de Koning et al., 2023) as can be observed in DTs of both marine ecosystems (Mészáros et al., 2024) and terrestrial systems such as grasslands (Taubert et al., 2024), crop production (Chalat et al., 2024), invasiveness of species (Khan et al., 2024), pollinators (Groeneveld et al., 2024), forest biodiversity dynamics (Afsar et al., 2024) and bird migration^1,2^ (Ovaskainen et al., 2024). Despite the demonstrated and wide ranging efforts of creating ecological DTs, no unified framework has been established governing structure and outlining common terminologies of DTs for ecological research and management (Schigel et al., 2024). Consequently, while the current implementation of DT frameworks in ecology remains in its early stages, it is crucial to establish clear concepts and unified frameworks to guide their development and prevent burgeoning fragmentation in terminology and design philosophies from becoming a significant barrier to progress. Deviations in implementations are only natural when new technology is adapted by a field of science, and oftentimes these deviations are good at this stage for exploring possibilities. However there are a range of technical challenges for ecological modellers that make ecological DTs unique. In the long-term, harmonising existing variation among ecological DT initiatives and rein in the potential for increasing confusion over the application of DTs to ecological applications, a unified well-structured design framework for ecological DTs is essential. This framework should encompass not only the technical aspects of data-model fusion but also ecological expertise, ensuring that the resulting DTs represent the intricacies of natural systems.

Conceptualisation of such a design framework encompassing the enormous variety of potential applications for DTs in ecological research/management is non-trivial as a host of research questions and considerations must be addressed. These include: Firstly, what should be the fundamental layers and components needed for building dynamic data-driven DTs in ecology? Given different scales and resulting management pathways for ecosystems, a design framework for ecological DTs must be malleable to account for the breadth of potential applications and resulting management and observation processes. It should also govern the conversion of existing static models to DTs. Secondly, how can DTs be structured in ecology? For example, the real-time data ingestion characterising industrial applications of DTs are usually not feasible for ecological research. Thirdly, how can ecological data from diverse sources be integrated into this framework seamlessly? Data products and types supporting ecological research are as diverse as the systems and processes studied by contemporary quantitative ecology. Interoperability of such disparate data poses a serious challenge to the establishment of DTs at scale (Reichman et al., 2011).

In addressing the aforementioned challenges and opportunities we propose a comprehensive design framework, called TwinEco, for dynamic data-driven DTs in ecology. To do so, we (1) develop a structured framework that outlines the essential layers and components involved in the creation of dynamic data-driven DTs for ecological systems, (2) investigate data integration and data/model fusion techniques that can accommodate diverse and dynamic data sources, including remote sensing, sensor networks, manually processed data and ecological monitoring data, (3) assess the potential of Dynamic Data-Driven Application Systems (DDDAS) as a system design approach to ecological DTs to automate system understanding, decision-making, mitigation, and management strategies, and (4) make future recommendations on what is needed to steer the process of digital twinning in ecological use cases. The resulting framework has the potential to leapfrog the way ecological systems are studied, monitored, and managed. Ultimately, this knowledge can guide DT development for conservation efforts, inform sustainable land use decisions, and contribute to the preservation of biodiversity and ecological balance.

## 3. TwinEco: The Framework

### 1. TwinEco Layers

In order to define concrete components that drive a DT, we first have to define the dynamics of the twin with its physical counterpart. The dynamics between a DT and its physical counterpart involve a complex interplay of data exchange, feedback loops, and synchronisation to achieve observation, understanding and action (Figure 1). All these aspects can be categorised as the “layers” of a DT. Each layer has its own individual mechanics, but there also exists mechanics of interaction between the layers. We will be discussing these mechanics as well. Similar methods of layering have already been adapted in existing DT approaches that do not necessarily follow any common framework, but share commonalities in the design patterns of the underlying systems (Buonocore et al., 2022; Fissore et al., 2023; Sougioultzogloua and Cook, 2023; Morlot et al., 2024).

#### Physical State, S

In the context of DTs, the concept of state refers to the representation of the current or past states of the Physical and DTs. The Physical State (*S*) encompasses all the relevant variables and parameters that describe the parameterized state of the physical asset at a given time in the Physical Twin (Malik et al., 2020) (Figure 2). These variables may include physical properties, environmental conditions, operational parameters, and any other factors that influence the behaviour of the physical asset. The Physical State changes over time (t) due to natural or unnatural events. At each time-step (t), the Physical State changes as well.

**Figure 2:**
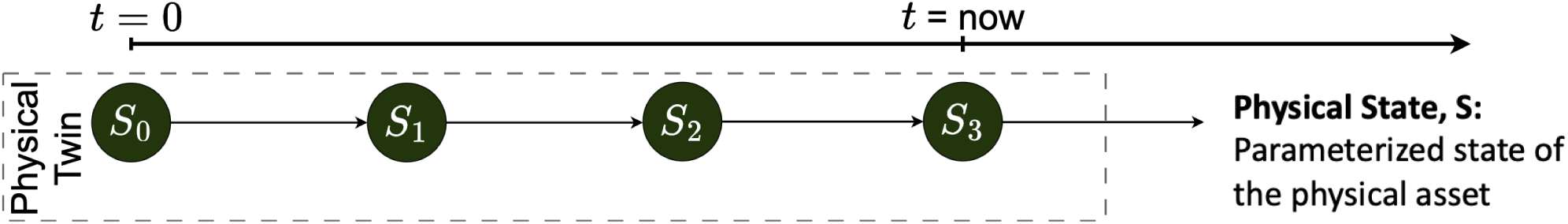
Parameterized state of the physical asset represented by the DT is defined as “Physical State” or S. S changes over time, starting at S_0_.

The Physical State is characterised by its complexity and dynamism, as it encompasses the ever-changing conditions and interactions within the physical environment. This complexity poses challenges for accurately capturing and representing the full extent of the Physical State in the DT. However, advancements in sensor technology, data analytics, and modelling techniques enable increasingly detailed and accurate representations of the physical asset (Onjaji et al., 2020).

#### Digital State, *D*

The Digital State (*D*) refers to the virtual representation of the physical twin within the DT (Malik et al., 2020). It comprises a set of variables and parameters that mirror those in the Physical State but are represented computationally. Like the Physical State, the Digital State also moves forward in time (*t*) and should be temporally in sync with the Physical State (Figure 3). The variables in the Digital State are typically captured through sensors, measurements, models or simulations and are used to simulate the behaviour of the physical system in the digital domain. This is essentially the “understanding” part of the DT.

**Figure 3:**
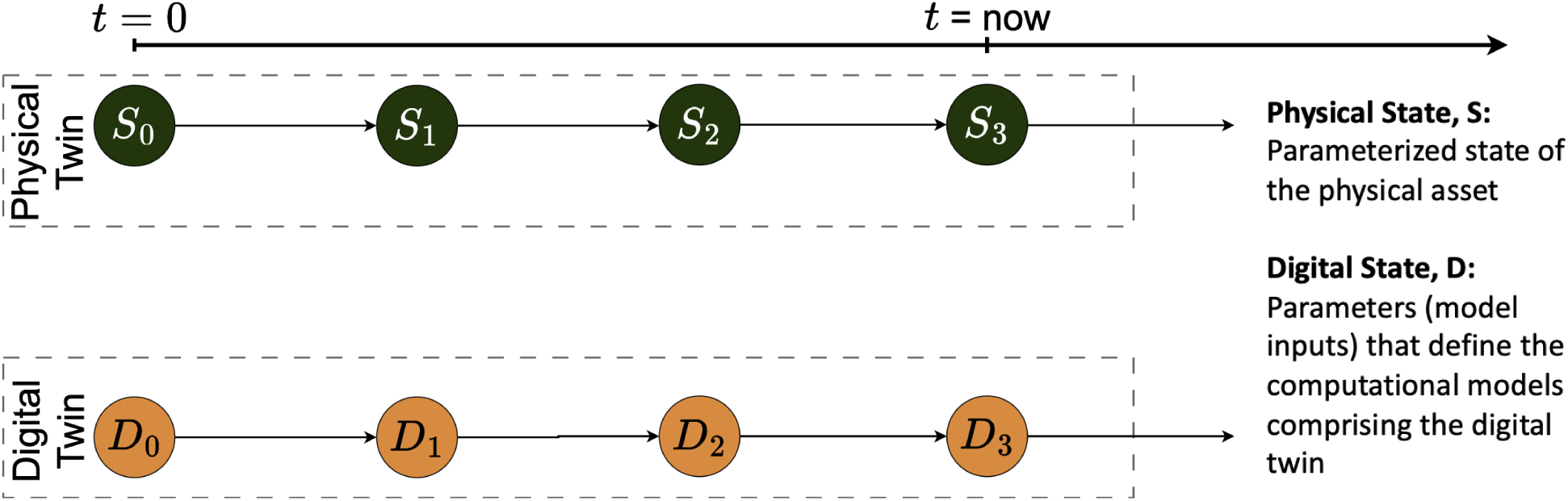
The Physical and Digital States represented in parallel, where the Digital State is a model or simulation that mimics the Physical State with a defined set of state variables, or “State Space” (DT component).

In contrast to the Physical State, the Digital State offers advantages such as controllability, reproducibility, and scalability (Wu and Li, 2021). Since it exists within the computational domain, the Digital State can be manipulated, analysed, and experimented with in ways that may be impractical or impossible in the physical realm. This flexibility allows researchers to explore different scenarios, optimise system performance, and predict future states of the physical twin (Wu and Li, 2022).

#### Observational Data, *O*

Observational Data (*O*) forms the link between layers *S* and *D* in Figure 4. The linkage between physical and digital states through observational data is fundamental to the operation of DTs, serving as the conduit through which real-world phenomena are mirrored and simulated in the digital realm (Malik et al., 2020). It is also important to note that *O* only captures a certain representation of *S* at a certain frequency and fidelity, therefore *O* should be seen as a simplified abstraction of *S*. Observational data, gathered from a variety of sources including field surveys, remote sensing, and monitoring stations, provide insights into the current state of the environment, encompassing factors such as biodiversity, habitat conditions, and ecosystem processes. These data serve as the backbone for DTs, acting as the primary means by which the digital representation of the ecosystem is updated and refined.

**Figure 4:**
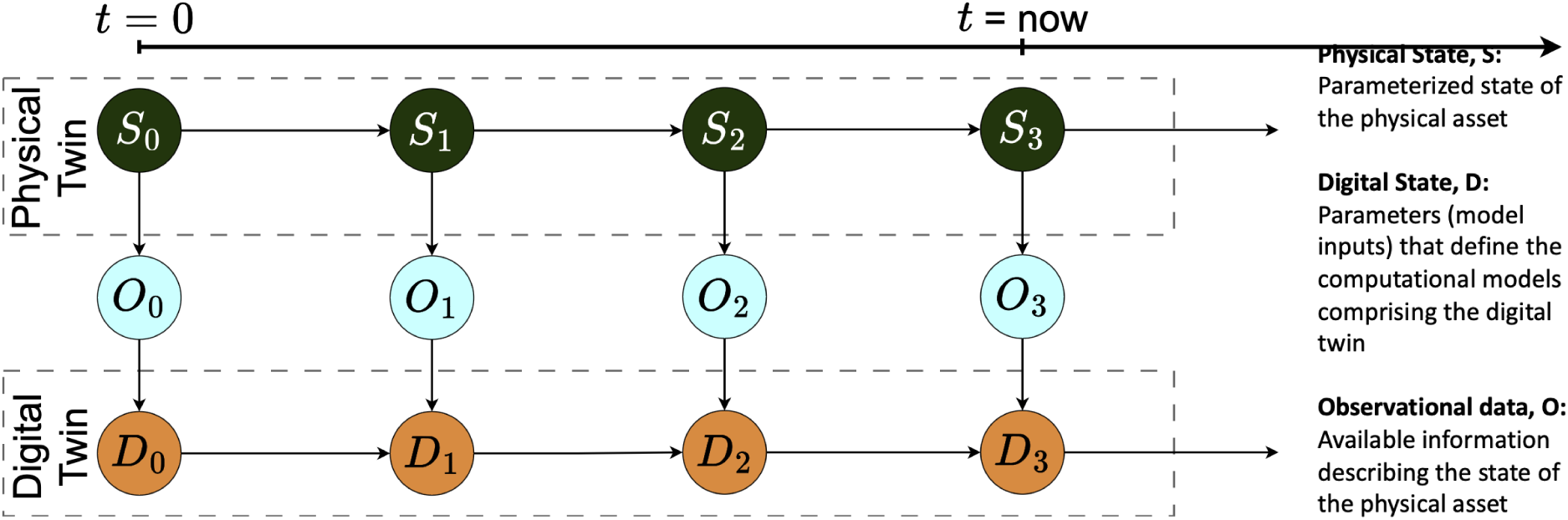
The Physical (S) and Digital (D) States are synchronised by the Observational Data (O), however the direction from S to D is not always linear as a temporal lag in O can shift the synchronisation. As S changes from S_0_ to S_n_, O_0_ and D_0_ also change to O_n_ and D_n_.

Moreover, observational data play a crucial role in validating the accuracy of ecological models embedded within the DT, ensuring that they capture the complexity and variability of real-world ecosystems. Through sophisticated data integration and analysis techniques, the DT synchronises its virtual state space with the ever-changing dynamics of the physical ecosystem. This synchronisation enables researchers to model and simulate ecological processes, predict future trends, and assess the potential impacts of human activities or environmental disturbances.

#### Adjusting Temporal Delay

There is a delay between when any changes happen in the Physical State, when any observation is made of the changes, and when the observation is added to the Digital State. To compensate for this delay, we introduce the concept of Temporal Delay.

Temporal Delay refers to the time difference between the timestamps of the Digital State (D_n_) and Physical State (S_n_). This can be represented as:

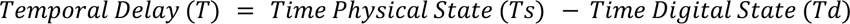

It can be seen that *T_d_* is completely dependent on when an observation is made (*T_o_*) and when the Physical State actually changes (*S_n_*) (Figure 5). For example, actual fish population size over time would be the physical state time series, incomplete and imprecise counts of fish sampled in a survey, or caught in a fishery, would be the observation time series, and the modelled fish count would be the Digital State time series. Such hierarchical time series structures have been previously described in State-Space and Non-Linear Dynamics modelling approaches, where the Temporal Delay has been referred to as the “hidden state” (Auger-Méthé et al., 2021).

**Figure 5:**
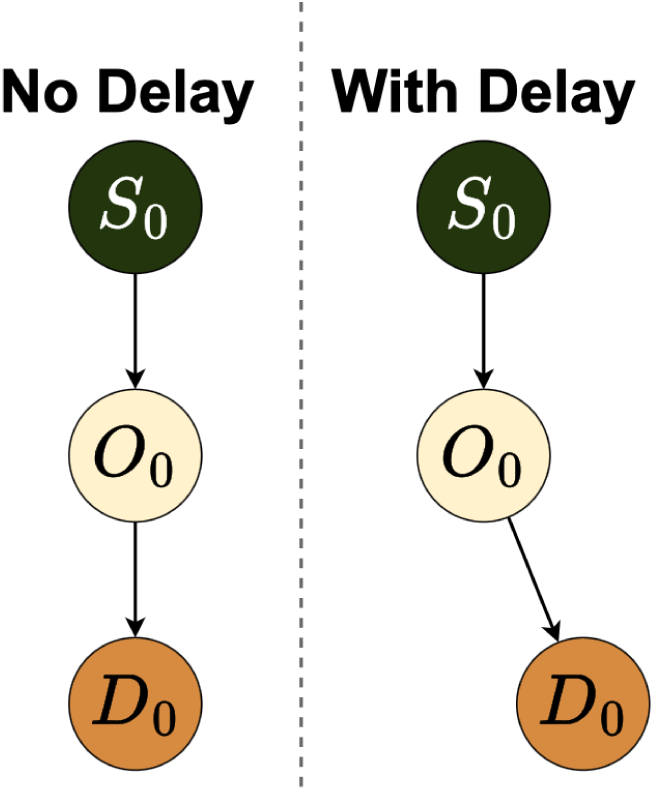
A visual representation of the Temporal Delay between when Physical Twin updates and when the DT updates. In many ecological cases, the Observational data representing a system is updated rather slowly, and usually considerable time has passed by the time models consume the data. Thus, the models represent a historic version of the Physical State.

Many aspects of ecology show stability over considerable spans of time. Hence, the temporal lag between Digital and Physical States may not matter in numerous cases. In contrast to engineering DTs, another differentiating feature of ecological DTs (and ecological data in particular) is that there are still many manual processes involved, both on providing the Observation Data as well as on the Control Inputs (managing the ecosystems based on insights from the DTs). Species range shift, time of flowering and other phenological aspects may not change overnight for example. Therefore, what matters perhaps is the acceptable length of temporal delay. The Temporal Delay is inherent in all DTs, though the degree of the lag varies. In most cases, an observation (*O*) is made of something (*S*) and its state is simulated (*D*). *O* and *D* cannot happen at the same time. The lag is between the observed (*O*) and predicted (*D*) states, not between the actual (*S*) and predicated (*D*) states.

#### Control Inputs, *U*

Control Inputs (*U*) in the context of DTs represent actions or interventions that can be applied to the physical system based on insights or predictions generated from its digital counterpart (Malik et al., 2020) (Figure 6). These inputs serve as a mechanism to influence and modify the Physical State of the asset in response to changes detected or anticipated in the Digital State. Control Inputs are a crucial component of DTs, enabling stakeholders to actively manage and optimise the behaviour of complex systems across various domains (Buonocore et al., 2022).

**Figure 6:**
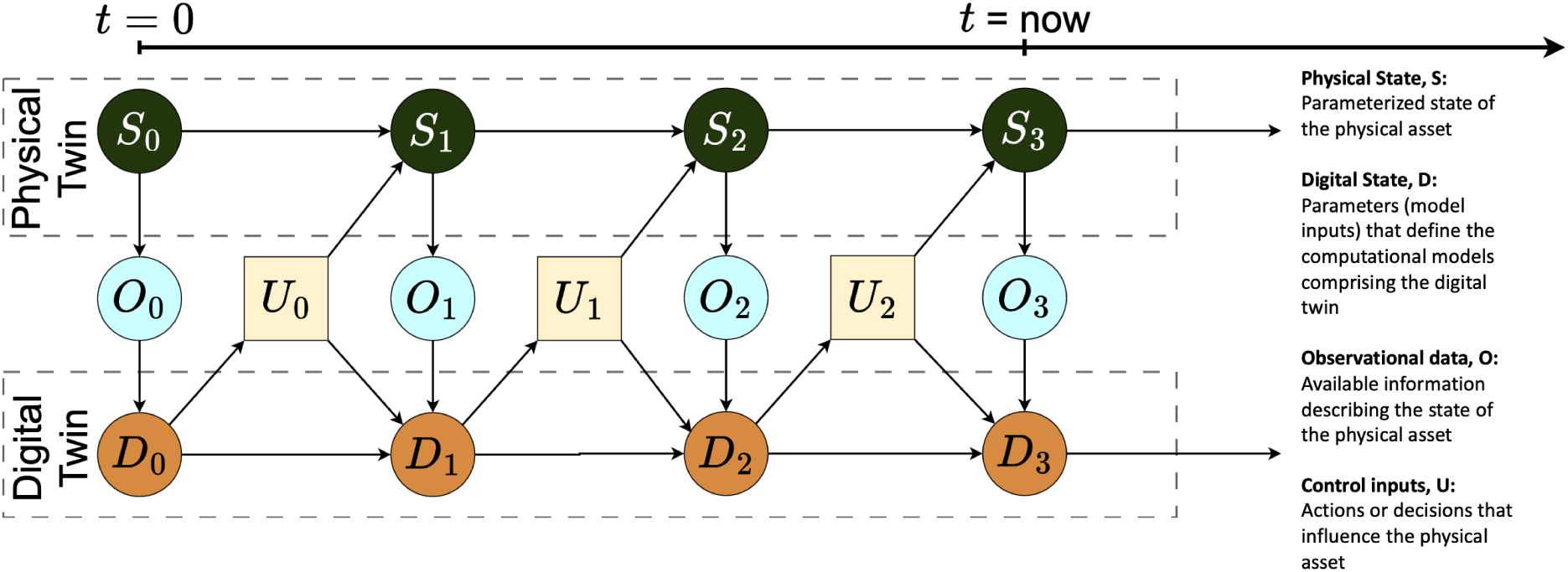
As the Digital State (D) updates at each time-step (t) based on new observational data, the computational modelled/simulated outputs can be used to drive a set of Control Input (U) at each step that changes the Physical State (S) of the asset.

In ecology, Control Inputs can either be of direct *<u>interventional</u>* type or of *<u>policy</u>* type. Policy controls inputs are basically decisions that can be made based on the DT output. Such Control Inputs can encompass a wide range of actions that can be applied to the physical asset, including adjustments to operational policy parameters, deployment and management of resources, and implementation of biodiversity control strategies. These actions are typically informed by analyses conducted within the DT, which may include optimization algorithms, predictive models, or decision-support systems. By leveraging the insights derived from the DT, stakeholders can identify optimal control strategies that maximise performance, efficiency, or other desired objectives while minimising risks or undesirable outcomes (Malik et al., 2020).

Here are examples of Interventional and Policy Control Input types:

##### 1. Interventional Control Inputs

In the management of a freshwater ecosystem such as a lake or reservoir using DTs, ecological Control Inputs could involve the manipulation of water flow rates (Qiu et al., 2023). By adjusting the flow rates of water into or out of the ecosystem, resource managers can influence factors such as water temperature, nutrient levels, and habitat availability. For instance, during periods of high nutrient runoff from surrounding agricultural areas, managers may increase the outflow rate to prevent eutrophication and maintain water quality. Conversely, during dry periods, managers may decrease the outflow rate to conserve water and maintain habitat integrity for aquatic species. These Control Inputs are informed by insights from the DT, which simulates the ecological dynamics of the system and predicts the impacts of different flow management strategies on ecosystem health and resilience (Qiu et al., 2023).

##### 2. Policy Control Inputs

A policy control input in ecology comprises suggested actions by the DT that help in implementing regulations or management strategies aimed at conserving endangered species or protecting critical habitats (Sharef et al., 2022). Such suggested actions or decisions from the DT, which may include predictive models of species distribution and population dynamics, can inform the development of policies that mitigate threats to biodiversity and ecosystem health (Scheibmeir and Malaiya, 2022). For instance, the DT could suggest establishing protected areas, habitat corridors, or zoning regulations based on projections of habitat loss due to urban development or climate change, helping policymakers safeguard key habitats and mitigate fragmentation. These policy interventions aim to address broader societal and environmental goals by integrating ecological knowledge with regulatory frameworks and stakeholder engagement, ultimately shaping land-use decisions and promoting sustainable development practices. In such cases, the policy could even be dynamic, and adjust with the changing DT outputs. It is worth mentioning that Policy Control Inputs add more time to the Temporal Delay, as implementations and subsequent changes take time to be implemented and show results.

#### Evaluation, *Ô*

Evaluation (*Ô*) of the Digital State based on Observational Data is a critical step in ensuring alignment between the virtual representation and the real-world conditions (Malik et al., 2020) (Figure 7). Discrepancies between the two can reveal areas where the DT may be inaccurately representing the physical twin or where Control Inputs may need adjustment. For example, if the DT predicts a decrease in species occurrence following the implementation of a control strategy but observational data show no corresponding reduction, it may indicate a need to refine the model or reassess the effectiveness of the control inputs.

**Figure 7:**
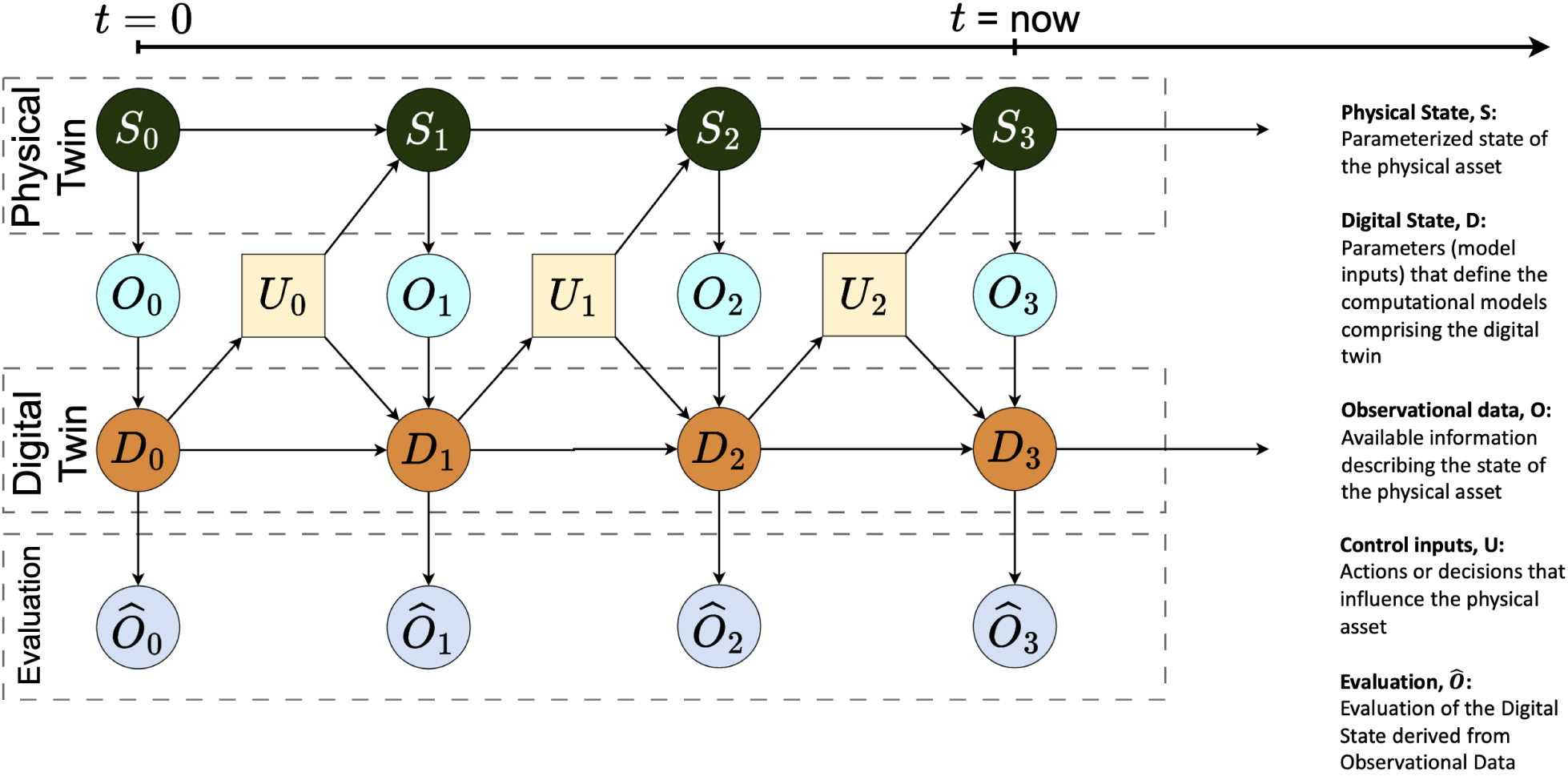
After the output of Digital State is updated after a model/simulation run, it is imperative to validate the results when compared to the Observational Data (O) in order to keep the Digital State (D) aligned with Physical State (S) at a desired fidelity and frequency.

Moreover, the evaluation of the Digital State serves as a feedback mechanism for validating the accuracy and reliability of the DT. By comparing simulated outputs to observed data, stakeholders can assess the fidelity of the DT in capturing the complex interactions and dynamics of the physical system. This validation process is essential for building trust and confidence in the DT as a decision-support tool for managing real-world systems (Trantas et al., 2023).

#### Rewards, *R*

Rewards (R) serve as evaluative metrics or objectives that guide the optimization of system behaviour towards an ideal state (Malik et al., 2020) (Figure 8). Rewards are typically defined based on the goals, objectives, or performance criteria established for the system being modelled. By quantifying the desirability or effectiveness of different Digital States or trajectories, rewards provide a feedback mechanism that informs the selection and application of Control Inputs to steer the system towards desired outcomes. Rewards can also be used to adjust the Temporal Delay if the Digital State is not at a desired fidelity or frequency.

**Figure 8:**
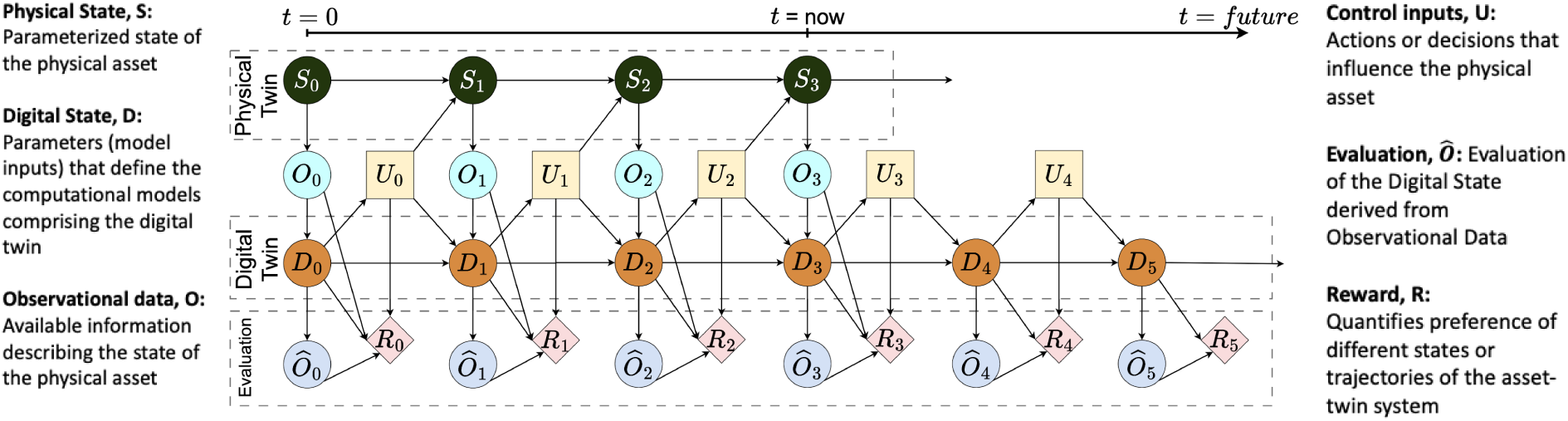
The complete overview of the layers of DTs in the TwinEco framework. Rewards (R) are linked to the Digital State (D), Observational Data (O), Evaluation (Ô) and Control Inputs (U).

Rewards are often used in conjunction with reinforcement learning algorithms, a type of machine learning approach that enables autonomous decision-making and control based on feedback from the environment where Q-learning is an algorithm that finds an optimal action-selection policy for any finite Markov decision process (MDP) (Dewey, 2014). In this context, rewards act as signals that reinforce or discourage specific actions taken by the DT in response to changes in the Physical or Digital States (Zekri et al., 2022). By associating positive rewards with actions that lead to desirable outcomes and negative rewards with actions that result in undesirable outcomes, DT algorithms can iteratively learn and optimise control strategies to achieve desired objectives.

For example, in the management of a water distribution network, rewards could be defined based on criteria such as system efficiency, water quality, and customer satisfaction (Zekri et al., 2022). By assigning higher rewards to Control Inputs that reduce water losses, improve water quality, and meet demand while minimising energy consumption, the DT can learn to optimise pump schedules, valve settings, and flow rates to achieve these objectives. Over time, the DT algorithm adjusts control strategies based on feedback from the environment, gradually steering the system towards an ideal state characterised by efficient water delivery and minimal waste.

In ecological applications, Rewards could be defined based on conservation goals such as biodiversity conservation, habitat restoration, or ecosystem resilience (Buonocore et al., 2022). For instance, in the management of a protected area, Rewards could be assigned to Control Inputs that enhance habitat quality, support endangered species, and mitigate threats such as invasive species or habitat degradation. By incentivizing actions that promote ecological health and resilience, Rewards provide a framework for prioritising management interventions and guiding decision-making in complex and dynamic ecosystems.

Overall, Rewards in DTs serve as a mechanism for aligning system behaviour with predefined objectives, enabling autonomous decision-making and ability to steer the physical asset towards desired states. By quantifying the desirability of different outcomes and providing feedback to learning algorithms, Rewards empower DTs to learn and adapt to changing conditions, ultimately facilitating the optimization of system performance and the achievement of strategic goals (Malik et al., 2020).

### 2. TwinEco Components

After defining the layers of DTs, the next step is to define the interaction between these layers. For this, certain components are suggested here. The components define the mechanics of the dynamics between the layers, meaning they realise the movement from *S* to *O* to *D* to *U* and so on. Inside a DT software environment, the components can be implemented in various software forms, allowing developers to tailor the system to their specific use cases and preferences. These components can be designed using Object-Oriented Programming (OOP) principles, where each component is represented as a class with its own attributes and methods, facilitating modularity and reuse. Alternatively, developers might employ functional programming techniques, encapsulating each component’s functionality within discrete functions that can be composed and reused across the system. In other scenarios, simple scripts might suffice, particularly for straightforward tasks or data processing pipelines. This flexibility of TwinEco in implementation ensures that DT environments can be customised to meet the diverse requirements of different ecological applications, whether they involve complex simulations, real-time data integration, or iterative model adjustments. By accommodating various programming paradigms and structures, DT systems can leverage the strengths of each approach, promoting efficient development, ease of maintenance, and scalability.

The Dynamic Data-Driven Application Systems (DDDAS) paradigm prioritises the integration of real-time data streams within computational models, facilitating adaptive decision-making and system control (Darema, 2004). Within the realm of DTs, DDDAS further extends this principle to develop dynamic, data-driven simulations that interact in real-time with evolving conditions within the physical system. By integrating feedback loops and state management mechanisms, this approach elevates the accuracy and adaptability of DTs (Malik et al., 2020). Consequently, within the TwinEco framework, individual components are constructed in the DT application to fulfil the functions of feedback loops and state management, effectively extending the capabilities of DDDAS (Table 1).

**Table 1:** This table encapsulates the essential components of the TwinEco framework, outlining their roles and interactions in creating a robust and flexible DT system for ecological applications. The type of components have been colour coded as red for State Management and green for Feedback loop components.

| Component | Type | Purpose |
| --- | --- | --- |
| State Space | State Management | The definition of input and output parameters of the DT. |
| Virtual Sensor | Feedback Loop | Detect changes in data sources and ensure new data availability before triggering other DT components, preventing data duplication. |
| Intaker | Feedback Loop | Pull new data into the system. |
| Processor | Feedback Loop | Clean and validate source data to fit desired rules and structures, ensuring data quality and consistency for the DT system. |
| Assimilator | Feedback Loop | Assimilate the new data into the existing/previous data and model, ensuring that the DT remains up-to-date with the latest information from the physical system. |
| Simulator | State Management | Run ecological models and simulate system dynamics based on data inputs and predefined parameters. |
| Actuator | Feedback Loop | Update the physical state based on the digital state, including ecological interventions and policy recommendations (Control Inputs). |
| Logger | Feedback Loop | Log events inside the DT to check whether all components function. |
| Indexer | State Management | Build a main time-line. Record data, modelling, and log files associated with each timestep, enabling stakeholders to track changes over time and examine different states of the DT. |
| Diff-Checker | State Management | Track changes between observational data and the digital state for each DT execution at each time-step, providing metadata for version control and system monitoring. |
| Evaluator | State Management | Validate models against empirical data to ensure accuracy and reliability, comparing simulated outputs with observed real-world conditions. |

#### Feedback Loop

In a DDDAS-enabled DT system, feedback loops play a crucial role in closing the loop between the physical and digital realms. These feedback loops enable real-time interactions between the DT and the physical system, facilitating the exchange of information, analysis, and control actions (Malik et al., 2020).

#### State Management

State management is another critical aspect of DDDAS-enabled DTs, ensuring that the digital representation accurately reflects the current state of the physical system. This involves maintaining a dynamic and up-to-date model of the system’s state, incorporating real-time data streams and adjusting model parameters as conditions evolve (Malik et al., 2020). State management mechanisms facilitate the integration and fusion of heterogeneous data sources, harmonising disparate data streams to create a cohesive and consistent view of the system’s behaviour.

#### State Space

Ecological systems exhibit complexity across broad temporal and spatial scales. Despite this complexity, ecological modelling approaches seek to distil the intricate dynamics of these systems into a set of manageable input and output variables. These variables collectively define the state of the system they represent and are collectively referred to as the State Space.

The State Space constitutes a fixed set of parameters consumed and outputted by the DT application. These parameters, once defined, evolve through time, shaping the dynamics of the DT. Moreover, the State Space establishes guidelines for the characteristics of the fixed parameters within the software, such as their data types (e.g., integer, float, string, date, etc.).

Ecological data is heterogeneous, stemming from diverse methodologies and standards employed for data collection in the field, alongside a lack of standardisation practices across studies (Niu et al., 2014). Consequently, a predefined State Space serves a crucial role in enabling the DT to manage this heterogeneity in its state data effectively. By imposing a structure, the State Space ensures that data adheres to desired formats, thereby facilitating the integration and harmonisation of disparate data sources to yield desired outcomes within the DT. In essence, the State Space functions akin to a schema or data model for the DT, guiding the organisation and interpretation of ecological data for effective modelling and analysis.

For instance, consider a DT modelling the dynamics of a forest ecosystem. One of the state variables within the State Space could represent the population density of a species like the red squirrel within a specific area of the forest. This state variable would capture the current population size of red squirrels in that area, providing valuable information about the health and dynamics of the ecosystem. As the DT evolves, this state variable would change dynamically in response to factors such as predation, habitat loss, or resource availability, reflecting the complex interactions within the ecosystem.

#### Virtual Sensor

Virtual Sensors (vSensors) serve as the initial point for triggering execution within the DT, detecting changes and updates in data sources across virtual spaces such as Application Program Interfaces (APIs), data files, and databases. They operate irrespective of whether these sources are directly connected to the DT (e.g. Internet of Things–IOT sensors), or obtained from third-party providers like research infrastructures. This ensures that other DT components are only triggered when new data is available for further processing, minimising data redundancy within the DT.

Change detection involves the DT actively monitoring the current state of data sources and comparing them for any alterations. For instance, a researcher utilising APIs of a research infrastructure to access species presence data can regularly query these APIs to ascertain the availability of new data on the servers. Virtual Sensors are tailored to each data source, accommodating unique formats, structures, and access protocols. Moreover, many data sources lack standardised metadata protocols, further complicating data integration processes.

In some cases, the DT first has to retrieve data by Intakers and subsequently check for changes by Virtual Sensors, necessitating periodic downloads and assessments to ensure that the data is up-to-date and accurate. Therefore, the order of which component comes first (vSensor or Intaker) depends on the source of the data and the available tools.

#### Intaker

Intakers are responsible for extracting raw observational data from diverse data sources. These sources encompass a broad spectrum of possibilities, including APIs, databases, file storage systems, repositories, and more. In addition to retrieving data, Intakers play a crucial role in validating the integrity and format of the data to ensure it aligns with the expectations of the DT.

For ecological applications, Intakers could be employed to download sensor data from various environmental monitoring stations scattered across a region. For instance, Inktakers might retrieve data from weather stations measuring temperature, humidity, and precipitation, or from soil moisture sensors deployed in agricultural fields. In another scenario, Intakers could pull data from satellite imagery repositories to track changes in land cover and vegetation density over time. These examples demonstrate the versatility of Intakers in sourcing data from disparate sources to fuel ecological modelling and analysis within the DT framework.

In many ecological use cases, Intakers must communicate with diverse research infrastructure APIs and file systems, complicating the data intake process. They handle various data formats and protocols, manage authentication and authorization, and integrate and harmonise data to fit the DT’s predefined state space. Additionally, they must ensure real-time data collection, implement robust error handling, and manage metadata for data integrity and traceability. For example, Intakers might download sensor data from cloud storage or interact with APIs, requiring them to navigate authentication, parsing, and data transformation challenges. These complexities underscore the need for sophisticated Intaker mechanisms to ensure reliable and accurate data ingestion into the DT system.

#### Processor

Processors play a critical role in the refinement and validation of source data within a DT system, ensuring adherence to desired rules and structures. This encompasses a spectrum of tasks, from addressing geospatial requirements such as transforming projections and Coordinate Reference Systems (CRSs), to harmonising taxonomic classifications. Furthermore, Processors are responsible for organising the data into specified structures, such as matrices, lists, or dataframes.

Processors also enforce data quality standards and ensure compliance with established data protocols within the DT. Given the automated nature of DT software, this step is important in preventing disruptions during data processing and analysis.

Data cleaning in ecological contexts presents a multifaceted challenge. Ecological datasets are often characterised by complexity and variability, requiring extensive filtering, cleaning, transforming, and validation procedures. Indeed, ecological models commonly find that the time invested in data cleaning far exceeds that allocated to subsequent analysis and modelling tasks. Automating this process poses significant challenges. Consequently, Processors emerge as a complex yet essential component of ecological DTs, serving as a linchpin for ensuring the integrity and reliability of data-driven analyses within ecological systems.

#### Assimilator

Assimilators assimilate the processed data into the existing data structure within the DT from previous DT runs. At this stage, the assimilator also handles tasks like versioning of the Observational Data, creating relations within the datasets (assigning unique identifiers), and storing the newly generated datasets into the storage system of the DT in the needed data format. The assimilators can also handle tasks related to cleaning the storage system by deleting the previous datasets or appending the new data to the previous dataset.

#### Simulator

The Simulator Component is the foundational element of a DT system, responsible for replicating the behaviour and dynamics of the physical system within the digital domain. It serves as the computational engine that drives the simulation of the physical system’s behaviour, enabling stakeholders to explore scenarios, predict outcomes, and optimise performance in a virtual environment. Simulators take the processed input datasets described in the State Space, and output the Digital State of the DT. A single DT could have a single or multiple Simulators.

At its core, the Simulator Component comprises mathematical or statistical models, algorithms, and computational techniques that capture the essential characteristics and interactions of the physical system. These models may range from simple mathematical approximations or statistical models to complex, process-based simulations, depending on the complexity and fidelity required for the specific ecological use case.

The Simulators take input data from various sources, including Observational Data, Control Inputs, and variables from the State Space, and use this information to simulate the behaviour of the physical system over time. By iteratively updating the state of the simulation based on these inputs, the Simulator Component generates a digital representation of the physical system’s behaviour, which can be visualised, analysed, and manipulated by stakeholders.

#### Actuator

In ecological modelling, direct intervention of models in the habitat or ecological process is not desired or possible. In many cases, in fact, the interventions are in the shape of policy recommendations and decisions. However, within the context of a DT system, predefined Control Inputs can guide the system towards an optimal state, driven by desired Rewards. Actuators serve as the components responsible for implementing these Control Inputs. With this in mind, Actuators can have two different types:

##### 1. Interventional Actuators

Interventional Actuators intervene directly in the Physical State to cause change. For instance, during periods of high nutrient runoff from surrounding agricultural areas, the Interventional Actuators may trigger an increase of the outflow rate to prevent eutrophication and maintain water quality.

##### 2. Policy Actuators

Policy Actuators, on the other hand, are action recommendations or decisions that steer the implementation of regulations or management strategies aimed at addressing ecological issues. These actuators facilitate the execution of policies designed to conserve endangered species or protect critical habitats. For instance, a predictive DT can inform Policy Actuators by providing insights into species distribution and population dynamics, thus guiding the development of effective policy parameters that safeguard biodiversity and ecosystem health.

A policy control input in ecology could involve implementing regulations or management strategies aimed at conserving endangered species or protecting critical habitats (Sharef et al., 2022). Insights from the DT, which may include predictive models of species distribution and population dynamics, can inform the development of policies that mitigate threats to biodiversity and ecosystem health (Scheibmeir and Malaiya, 2022).

#### Logger

Loggers serve as a vital tool for monitoring and recording events occurring within the software environment. Its primary function is to track the behaviour and interactions of various components within the DT to ensure that they are functioning as intended. By logging events and activities, Loggers provide valuable insights into the performance, reliability, and overall functioning of the DT system.

The Logger Component operates by capturing and storing information about key events, such as the execution of Control Inputs, updates to the Digital State, and responses from Actuators. This information is typically recorded in a structured format, including timestamps, event descriptions, and relevant contextual metadata. By maintaining a comprehensive log of events, the Logger Component enables stakeholders to trace the sequence of code executions and identify any logical errors in the algorithms or bugs in the software.

Loggers play a crucial role in system diagnostics and troubleshooting. In the event of errors or malfunctions within the DT, the log data can be analysed to pinpoint the root cause of the issue and facilitate corrective actions. By correlating events and identifying patterns in the log data, engineers and developers can gain valuable insights into the underlying dynamics of the DT system and implement improvements to enhance its performance and reliability.

#### Indexer

Indexers are a temporal indexing mechanism implemented within the TwinEco DT framework that serve as a crucial tool for maintaining a comprehensive record of data, Digital State, and Physical State evolution over time. By tracking timestamps and associating them with relevant data and files, this metadata infrastructure facilitates version control and enables stakeholders to navigate through different temporal states of the DT.

For instance, as the DT progresses from its initial state at *t*=0 to subsequent time-steps, the indexers diligently record all associated data, modelling outputs, and log files. This temporal labelling allows users to seamlessly traverse through various time-steps of the DT, whether it’s for retrospective analysis, model evaluation, or performance assessment.

The temporal indexing system finds applications in meta-modelling endeavours and evaluating model performance. By indexing single updates of the Digital State and correlating them with the timestamps of corresponding Observational Data recordings in the physical system, stakeholders gain insights into the temporal alignment between simulated and observed states. Three distinct types of Indexers fulfil this function: Observational Data indexers, Digital State indexers, and Physical State indexers. These Indexers provide answers to critical questions such as when the data was recorded, when the prediction was produced or the DT was updated, and what specific time and date the DT predictions represent. This comprehensive temporal indexing infrastructure enhances the utility and transparency of DT systems, empowering stakeholders to make informed decisions and derive valuable insights from temporal data dynamics.

#### Diff-checker

Diff-checkers serve a pivotal function within the TwinEco framework by monitoring alterations in both Observational Data and the Digital State during each DT update. These mechanisms, essentially metadata, provide an invaluable record of what changes occur over time.

The utility of Diff-checkers extends across various domains, including metamodelling, debugging, model evaluation, and decision-making processes. Diff-checkers offer comprehensive insights into system dynamics, enabling stakeholders to track changes not only from a data perspective but also from a model viewpoint, thereby furnishing a holistic understanding of system behaviour. In essence, Diff-checkers play a role in monitoring system dynamics, facilitating informed decision-making and enhancing the value proposition of DTs over traditional approaches.

For instance, consider a DT designed to model the distribution of a particular species within a forest ecosystem. The Diff-checker would continuously compare updates in Observational Data, such as species occurrence records from field surveys or remote sensing data, with the predictions generated by the DT, capturing what exactly changes from one time-step to the next. If the DT predicts an increase in the population density of the species within a certain area, but the Observational Data indicate a decline or no change, the Diff-checker could flag this discrepancy. This could prompt further investigation into potential factors influencing the discrepancy, such as habitat degradation, climate change effects, or inaccuracies in the model assumptions. Overall, the Diff-checker serves as a valuable tool for ensuring the accuracy and reliability of the State Space and the Simulators within DT systems.

#### Evaluator

The role of Evaluators within the development and validation of DT systems is paramount. The Evaluator Component assumes a pivotal role in the pre-deployment phase, serving to validate the accuracy and reliability of the models against empirical data. This validation process entails rigorous comparison of simulated outputs with observed behaviour in real-world conditions. Through iterative refinement, the models are honed until they faithfully replicate the behaviour of the physical system at the desired frequency and fidelity.

The nature of evaluation undertaken is contingent upon the type of Simulator utilised within the DT framework.

#### Mapping components to layers

As shown previously, the TwinEco framework is organised into distinct layers, each comprising specific components that collectively enable comprehensive ecological modelling. The components introduced above can be mapped to the layers of the TwinEco framework to create the interactions between layers (Figure 8).

As the DT transitions from the Physical State to the Digital State at each time-step, it incorporates components such as Virtual Sensors and Intakers, which are responsible for gathering and integrating raw data from various sources. This layer ensures that the DT has access to up-to-date and relevant Observational Data.

In the Observational Data layer, components like Processors and Diff-Checkers play a crucial role in cleaning, validating, and tracking changes in the data from one execution to the next. This layer transforms raw data into structured formats suitable for modelling and analysis. The Digital State layer includes the Simulator, which runs the ecological models, and Actuators, which implement control inputs based on model predictions.

The Evaluation layer comprises Evaluators and Indexers. Evaluators validate the accuracy and reliability of the models by comparing simulated outputs with empirical data, while Indexers manage metadata and ensure version control, allowing for the tracking of changes over time. Together, these layers and components create a robust and adaptable framework for developing DTs in ecology. Figure 9 illustrates the mapping of these layers to their respective components as single time-steps, providing a visual representation of the TwinEco framework.

**Figure 9:**
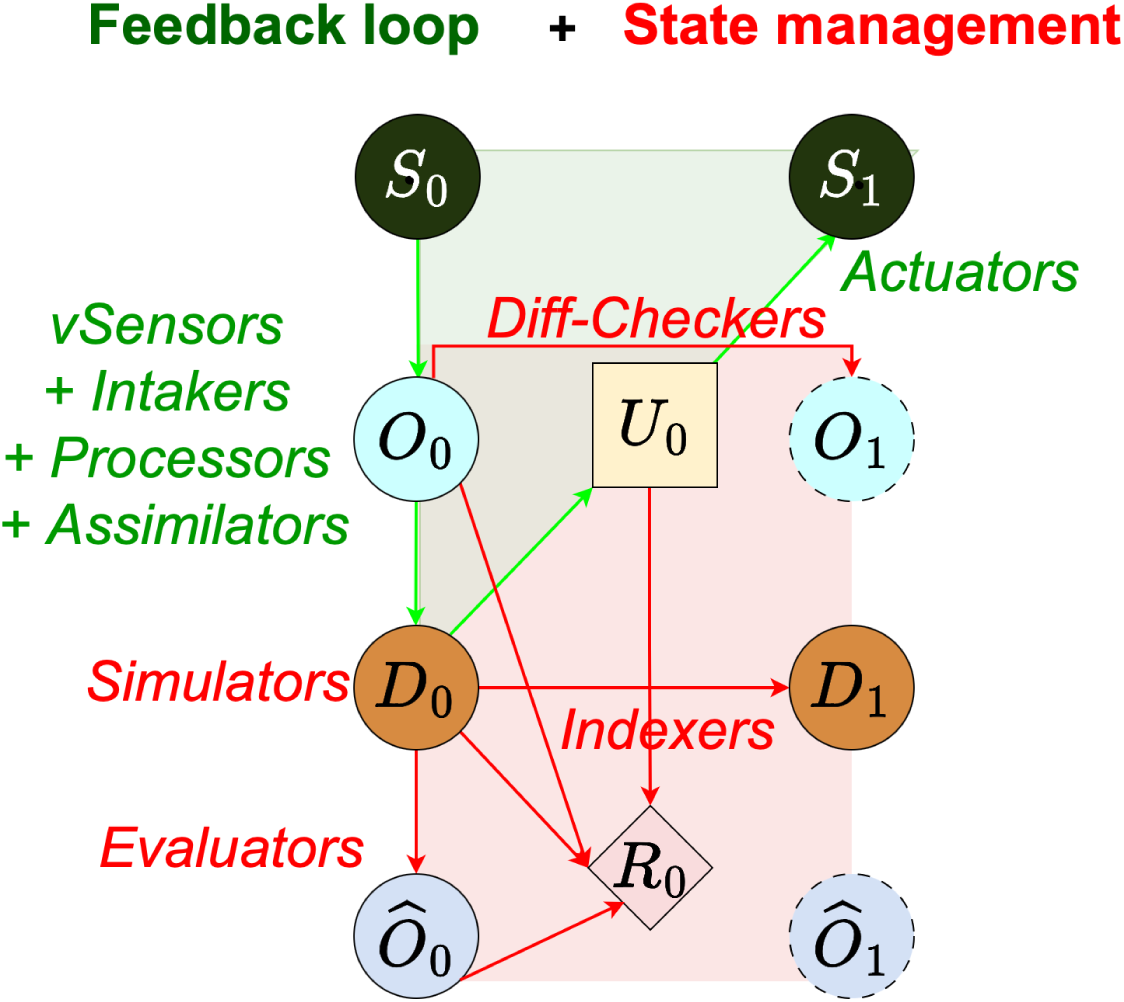
Mapping of the components to the layers of the TwinEco framework in a single timestep, where feedback loop (green) and state management (red) components are colour coded.

## 4. Ecological Relevance and Future Recommendations

### Applicability and Computational Representation in Ecology

The flexibility and modularity of the TwinEco framework make it an ideal tool for addressing the broad range of digital twinning use cases in ecology, each with unique data requirements and modelling complexities. Ecosystems are often characterised by diverse processes — ranging from species interactions to biogeochemical cycles — that occur across different spatial and temporal scales. As such, designs of ecological DTs need to be highly adaptable, able to integrate multiple data streams, and capable of responding dynamically to environmental changes. Table 2 demonstrates how TwinEco supports these demands through its comprehensive components. These components can be implemented computationally in various ways depending on the use case requirements and developers’ choices.

**Table 2.**
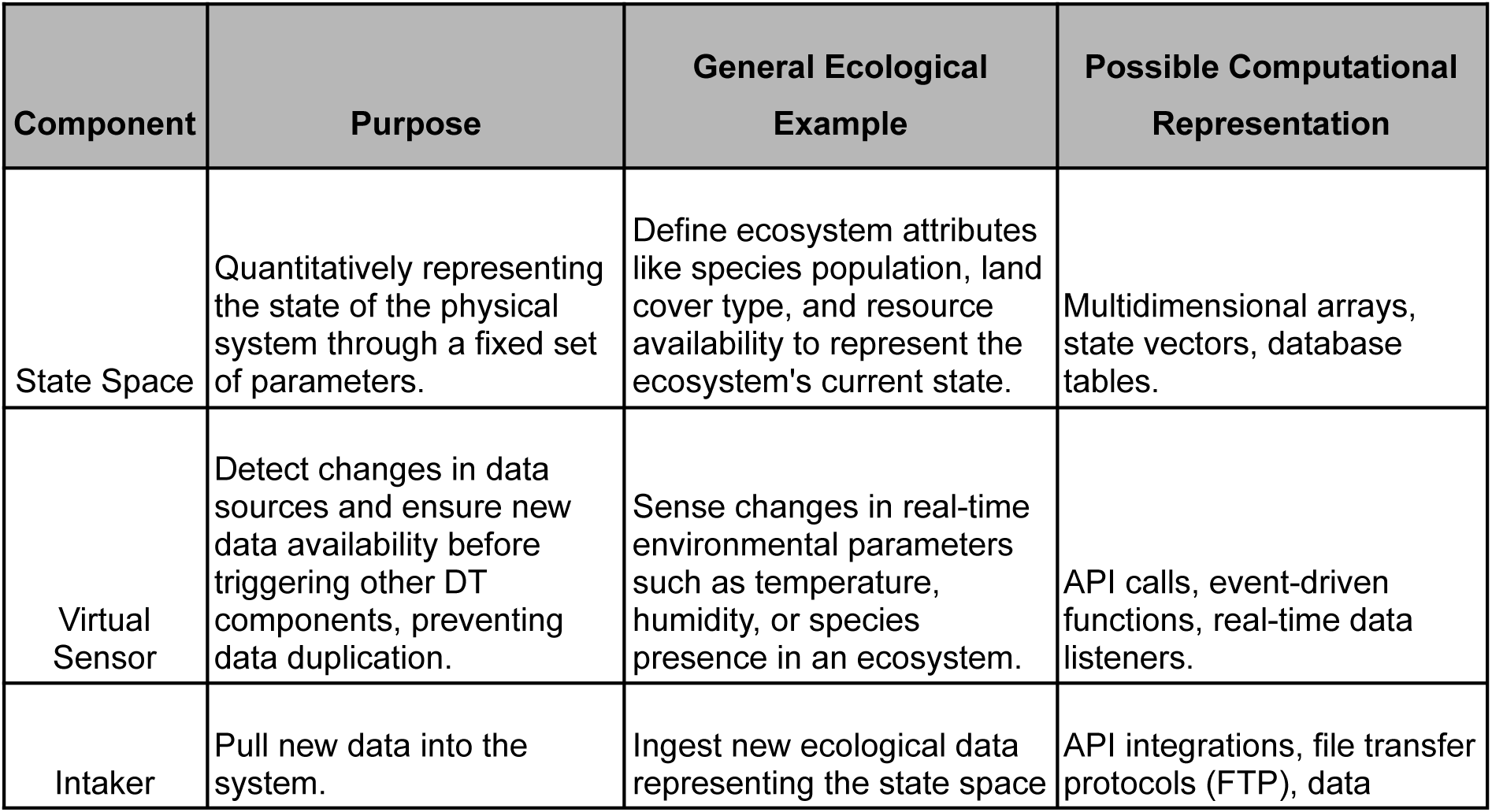
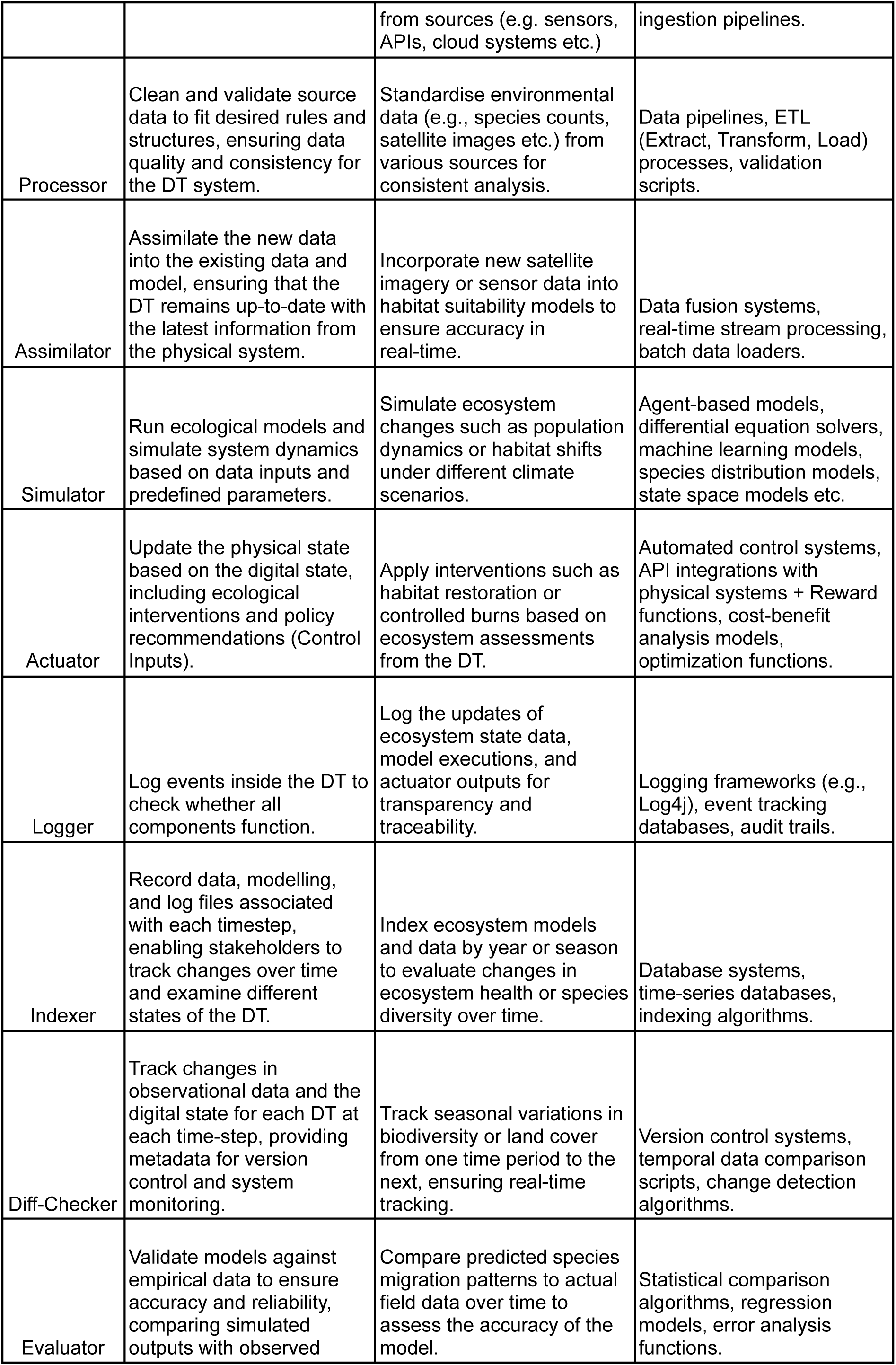
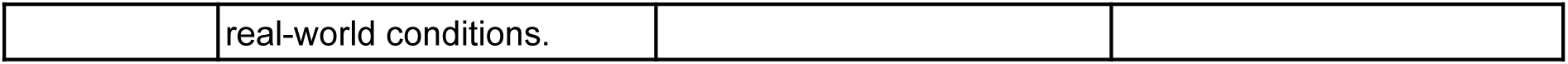
Key components of the TwinEco framework with ecological examples and computational representations. Each component of the framework is tailored to address specific aspects of ecological digital twinning, from real-time data intake and processing to model simulations, decision-making, and actuation. The examples focus on a forest ecosystem, highlighting applications like monitoring tree health, simulating growth dynamics, and optimizing management interventions. Computational representations and software implementation examples (e.g., functions, scripts, OOP) demonstrate the adaptability and practicality of each component in real-world ecological and computational contexts.

Ecological use cases are broad, encompassing not just terrestrial systems like forests and grasslands but also aquatic ecosystems, urban landscapes, agricultural environments, and even microbial communities. The framework’s focus is broad enough to handle different data types (e.g., sensor data, remote sensing, field surveys, or citizen science data), assimilate real-time inputs, and validate model outputs through empirical data comparison (Jorgensen & Bendoricchio, 2001). For instance, in urban ecosystems, where rapid anthropogenic changes impact biodiversity, DTs built following the design principles presented in TwinEco could integrate data from urban sensors to monitor air and water quality, providing insights into how urban expansion affects local species and ecosystem services. In contrast, in agricultural landscapes, DTs following our design specifications could predict crop yields or soil health, advising on sustainable farming practices in the face of fluctuating environmental conditions (Wu et al., 2006). Ultimately, the TwinEco framework can be applied to DT applications across ecological research disciplines due to its agnostic perspective on study systems and data sources, but consistent yet flexible design components.

However, while TwinEco is versatile, there may be certain ecological use cases where the framework’s structure may not fully apply. For example, ecosystems where the dynamics are predominantly stochastic and difficult to model — such as highly ephemeral or unpredictable systems like desert flash floods — might pose challenges for TwinEco’s more deterministic approach. In such cases, the reward functions or control inputs could struggle to define ideal states or interventions due to the inherent unpredictability of the system (Grimm et al., 2005). Moreover, systems where data availability is scarce, such as remote or deep-sea ecosystems, may hinder the effective use of Virtual Sensors or Simulators, given the need for real-time data feeds to support the twin’s continuous updating process.

Nonetheless, for most ecological use cases, TwinEco is a robust and adaptable framework for building DTs. It offers the capacity to integrate a variety of data sources, simulate complex interactions over time, and provide decision support for ecosystem management. For example, in forests, the TwinEco framework could help to develop DTs that monitor seasonal growth dynamics and predict responses to climate change, while in wetlands, DTs could track water quality and hydrological changes to ensure optimal habitat conditions for wildlife (Landi et al., 2018). Furthermore, the ability to use or omit certain components — like Actuators in ecosystems where direct intervention is not possible — enhances its applicability across a wide range of ecological applications, ensuring that the framework does not impose unnecessary components on simpler use cases. This flexible and modular design allows researchers to customise implementations based on their specific needs, using a range of programming techniques such as scripts, functions, and Object-Oriented Programming (OOP).

#### Future Recommendations

DTs herald a shift in ecological modelling, prompting rapid adoption for scientific research. With burgeoning interest among modellers, however, there arises a heightened risk of fragmentation in defining and structuring DT applications. This fragmentation could lead to divergent interpretations across communities, resulting in relevance confined within specific domains but diminishing outside reach. This challenge becomes particularly pronounced with initiatives like Destination Earth, aiming to construct comprehensive DTs of Earth and its myriad systems (Nativi et al., 2021). Consequently, the imperative for a unified DT framework, such as TwinEco, becomes more pressing than ever, ensuring coherence amidst the proliferation of diverse use cases.

**Recommendation 1:** Researchers building new DTs should be mindful of this risk of fragmentation of practices and aim to follow shared principles including frameworks such as TwinEco.

A DT can serve numerous purposes in ecology and life sciences. For example, satellite image based DTs can detect forest disturbances, map and monitor crop stresses, such as drought and disease, as well as crop maturity in near real time, providing alerts for necessary actions. Furthermore, DTs can access molecular data of medically important viruses to identify genetic changes, evaluate the potential for vaccine resistance, and inform measures to prevent the spread to other regions (Baaden, 2022). They can also detect changes in data availability to build and update habitat suitability models for species in real time, and enable real-time detection of invasive species spread.

In general, DTs in ecology are used to access and integrate data from different sources to calibrate models and produce the necessary knowledge in real time. Thus, in most cases, they are more about informing the state of knowledge regarding certain aspects of the model targets (the physical states) than about the state of the physical twin itself.

Depending on the specific aspect of an ecosystem or the taxa being modelled, DTs offer flexibility in the components and layers that could be used. For example, we may not need the evaluation layer/component to detect changes in molecular makeup of a virus. One can use DTs to update data on certain aspects of an ecosystem/taxa and inform the gap in sampling to advise where to sample next. In this specific case, for example, the “Diff-checker” component of the DT may not be that important. Additionally, functionality of multiple components could also be combined into certain use cases.

However, adhering to a common framework is crucial for making DTs interoperable or capable of communicating with one another. This common framework allows ecological DTs to integrate easily into initiatives like Destination Earth (Le Moigne, 2022; Ossing et al., 2023; Le Moigne 2024). It ensures they can use explanatory data from the same sources of the destination earth data lake and guarantees their sustainable existence and functionality. The components of DTs could be shared as “common” components by other twins, especially the components that pull and process data from common research infrastructure in ecology.

**Recommendation 2:** DTs in ecology should aim to follow common design patterns to increase interoperability of the twins across domains and for the twins to be able to reuse parts of each other to create newer, more novel DTs.

Unlike many traditional ecological modelling frameworks, establishment of an operational and useful analysis workflow using the TwinEco framework does not necessitate deployment of all components we have introduced here. This particular aspect of flexibility/modularity inherent to TwinEco presents two distinct opportunities as well as one marked challenge in adopting its principles into ecological research applications. Firstly, not requiring deployment of all structural components to achieve functionality makes TwinEco adaptable to many ecological applications. For example, many, if not most, ecological disciplines focus on the study of systems which are difficult to influence via actuator-driven feedback loops and thus will require study via TwinEco DTs without the actuator component. Secondly, clear communication of implementation of TwinEco components in ecological analyses pipelines can serve as a benchmark for quantification of degree to which a fully integrated DT has been established. This may serve as a guiding principle when designing studies as well as augment past research efforts by adding classifications to their workflows. Lastly, however, this modularity also begs the question “which components are required for a workflow to be called a digital twin?”. This is a non-trivial topic with potentially far-reaching consequences that ought to be discussed with the wider ecological research community (Ossing et al., 2023). TwinEco lays the foundation for such consensus-finding on what constitutes an ecological DT creating room for direct discussion of components, layers, and time-sensitivity of digital twinning applications.

**Recommendation 3:** Adjust frameworks like TwinEco to specific use cases, as not all components of the framework will be applicable to every DT.

DTs require dynamic real time input data while common data collection protocols in ecology result in a time lag and batch updated data sources. Recent experiments with real-time data collecting sensors point towards the need for new data streams more fit for use by DTs. However, automatic real-time data fusion across distributed networks of sensors add new even stricter requirements for machine-actionable and machine-interoperable data formats and semantics. On the other side, solving these operational challenges facilitates the up-scaling of monitoring of ecological systems.

**Recommendation 4:** Researchers building new monitoring systems should be mindful of enabling true machine interoperability and machine data fusion to support emerging DT systems.

Building DTs in ecology requires stable data streams facilitated by permanent research infrastructures. Current research infrastructures for biodiversity and ecology data in Europe were designed and implemented before the emerging implementation of DTs in ecology. Without the strong demand for real-time sensor data streams the current research infrastructures have been implemented with a time lag in data delivery (Li et al., 2023).

**Recommendation 5:** Research infrastructures for biodiversity and ecology data should aim to position themselves to become more fit for use by the emerging DTs and establish functionality and new data stream services for more real-time data delivery, and to work together with downstream data sources to mobilise such real-time data streams.

Digital twinning is new to ecologists and there is a need to establish a shared terminology and understanding of DT components and concepts (Korenhof et al., 2021). This paper introduces terminology to promote a shared community definition of components needed for building DTs in ecology. The interTwin project^3^ is working on establishing a shared vocabulary of terminology for key concepts of digital twinning.

**Recommendation 6:** Digital twinning engineers in ecology should contribute to and follow the emerging shared terminologies, conceptualisations, and component frameworks (such as the TwinEco proposed here).

Using the TwinEco framework, ecological dynamics can be observed and understood with continuous updates to models/digital representations of real world systems/processes, and in return the twins can put back actions/decisions in the real-world. The framework highlights the components that are needed to be able to do that. This allows modellers to convert existing static ecological models into DTs. The framework also helps to improve the understanding of the components and tasks that are involved in developing DTs. The framework explicitly models knowledge and processes/states over time, allowing for a more dynamic and realistic representation of ecological systems that updates as the ecological systems change. By integrating vast amounts of diverse data from various sources, the framework ensures a comprehensive and up-to-date understanding of ecological change.

## 5. Conclusion

The development and implementation of DTs in ecology mark a transformative advancement in ecological modelling. By integrating the Dynamic Data-Driven Application Systems (DDDAS) paradigm, the TwinEco framework offers a unified approach to constructing DT models that are adaptive, responsive, and capable of real-time data integration. This framework addresses the challenges of fragmentation in ecological DTs by providing an overview of the important layers and components of DTs, thereby enhancing clarity on the concept for ecological modellers and ensuring consistency in the basic architecture of ecological DTs (Boyes and Watson, 2022). Consequently, the TwinEco framework facilitates the creation of robust and reliable ecological models that can inform decisions or actions with unprecedented accuracy and timeliness.

A key strength of the TwinEco framework lies in its flexibility and modularity. This allows researchers to tailor DT components to specific ecological contexts, whether it involves tracking forest disturbances, monitoring crop health, or assessing the spread of invasive species. By enabling dynamic data processing and continuous model refinement, TwinEco helps ecological modellers to conceptualise complex ecological dynamics and respond to environmental changes swiftly. Moreover, the framework’s emphasis on interoperability ensures that DTs can seamlessly integrate with larger initiatives like Destination Earth, leveraging shared data sources and contributing to a comprehensive understanding of global ecological systems (Rao et al., 2023).

Future research and development in ecological DTs should focus on enhancing real-time data collection and integration capabilities of the Research Infrastructures, particularly through focusing on data streaming technologies and data standardisation. Efforts should be made to improve the interoperability of data sources, ensuring seamless communication and data exchange between different DT systems and across various ecological domains. Additionally, it is essential to demonstrate the framework’s utility through additional case studies. Documenting diverse applications will illustrate the TwinEco framework’s versatility in modelling various ecological systems and addressing a wide range of ecological challenges. These case studies will serve as practical examples showcasing the framework’s strengths and areas for improvement. Collaboration among ecologists, data scientists, and technologists will be crucial in driving these innovations forward. Establishing shared terminologies, best practices, and guidelines will also be essential to foster a cohesive community approach and to maximise the impact and utility of DTs in ecological research and management (Korenhof et al., 2021).

In conclusion, the TwinEco framework represents a step forward in understanding DTs in ecological modelling, providing a unified yet flexible approach to developing DTs. It addresses the critical need for data update integration and model adaptability, offering a powerful tool for advancing ecological research and management. As the field of ecological modelling continues to uptake cutting-edge technology, the adoption of frameworks like TwinEco will be essential in ensuring that DTs remain relevant, reliable, and capable of addressing the complex challenges facing our ecosystems today and in the future. By fostering collaboration and standardisation, TwinEco paves the way for more effective and coordinated efforts in ecological conservation and sustainability.

## 6. Acknowledgements

This study has partially received funding from the European Union’s Horizon Europe research and innovation programme under grant agreement No 101057437 (BioDT project, https://doi.org/10.3030/101057437). Views and opinions expressed are those of the authors only and do not necessarily reflect those of the European Union or the European Commission. Neither the European Union nor the European Commission can be held responsible for them. We acknowledge the EuroHPC Joint Undertaking for awarding this project access to the EuroHPC supercomputer LUMI, hosted by CSC^4^ (Finland) and the LUMI consortium through a EuroHPC Development Access call. We would also like to extend our gratitude to <u>Marija Milanović</u> for her invaluable assistance in designing and creating Figure 1.

## 7. Author Contributor Roles

Table 2 shows the contribution roles that have been ascertained using CRediT^5^ (Contributor Roles Taxonomy).

**Table 3:** The CRediT roles for the authors of this manuscript.

| Author | Roles |
| --- | --- |
| Taimur Khan | Conceptualization, Investigation, Methodology, Project administration, Supervision, Validation, Writing – original draft |
| Koen de Koning | Conceptualization, Writing – review & editing, Methodology, Validation |
| Dag Endresen | Conceptualization, Writing – review & editing, Validation |
| Desalegn Chala | Conceptualization, Writing – review & editing, Validation |
| Erik Kusch | Conceptualization, Writing – original draft, Writing – review & editing, Methodology, Validation |

<sup>4</sup> https://www.csc.fi
<sup>5</sup> https://credit.niso.org/

## 8. Declaration of generative AI and AI-assisted technologies in the writing process

During the preparation of this work the authors used ChatGPT in order to improve readability and grammar for sentences that were too long or awkward, as none of the authors are native English speakers. No aspect of this manuscript used ChatGPT for content, logic or reasoning. After using this tool/service, the authors reviewed and edited the content as needed and take full responsibility for the content of the publication.

## Footnotes

1 https://sensingclues.org/craneradar

2 https://biodt.eu/use-cases

3 https://github.com/interTwin-eu/dtc-glossary

4 https://www.csc.fi

5 https://credit.niso.org/

## Notes

### Competing Interest Statement

The authors have declared no competing interest.

### Summary of Updates

The arguments and the link to ecology are much stronger than the last version. I also took the time to develop a new figure, Figure 1, together with my colleague Marija and our co-author Erik. This new figure introduces the ideas much more cleaner and also introduce our figure-system in TwinEco in a lighter way. In that sense the build up is much cleaner now. We also heavily reworked the abstract, introduction, and discussion sections.

